# USP7 couples DNA replication termination to mitotic entry

**DOI:** 10.1101/305318

**Authors:** Antonio Galarreta, Emilio Lecona, Pablo Valledor, Patricia Ubieto, Vanesa Lafarga, Julia Specks, Oscar Fernandez-Capetillo

## Abstract

To ensure a faithful segregation of chromosomes, DNA must be fully replicated before mitotic entry. However, how cells sense the completion of DNA replication and to what extent this is linked to the activation of the mitotic machinery remains poorly understood. We previously showed that USP7 is a replisome-associated deubiquitinase with an essential role in DNA replication. Here, we reveal that USP7 inhibition leads to the ubiquitination of MCM7, a hallmark of DNA replication termination. In addition, USP7 inhibition leads to the ubiquitination of additional replisome components such as POLD1, which are displaced from replisomes. Surprisingly, this premature termination of DNA replication occurs concomitant to a generalized activation of CDK1 throughout the entire cell cycle, which impairs chromosome segregation and is toxic for mammalian cells. Accordingly, the toxicity of USP7 inhibitors is alleviated by CDK1 inhibition. Our work sheds light into the mechanism of action of USP7 inhibitors and provides evidence to the concept that DNA replication termination is coupled to the activation of the mitotic program.

## INTRODUCTION

The eukaryotic cell cycle is governed by cyclin/CDK complexes that coordinate an orderly transition between the different cell cycle stages (G1, S, G2 and M). As noted early on, among the different transitions, limiting mitotic entry before DNA replication is completed is of particular relevance (Enoch & Nurse, 1991, Hartwell & Weinert, 1989). Besides the fact that unreplicated loci physically impair chromosome segregation, the premature activation of mitotic kinases in S-phase leads to chromosome pulverization and cell death (Aarts, Sharpe et al., 2012, Duda, Arter et al., 2016). Likewise, early works revealed that the fusion of a mitotic cell with an interphase cell triggers premature chromosome condensation (PCC) in the later (Johnson & Rao, 1970). Consistent with the key role of CDK1 in the G2/M transition, premature mitotic entry can be induced by genetic or chemical strategies that increase CDK1 activity, such as wee1 deletion in fission yeast (Russell & Nurse, 1987), or the use of WEE1, ATR or phosphatase inhibitors in mammalian cells (Aarts et al., 2012, Ajiro, Yoda et al., 1996, Ruiz, Mayor-Ruiz et al., 2016). In this context, the main function of an S/M checkpoint should be to restrict CDK1 activity until DNA replication ends. However, while the presence of such a checkpoint has been long thought of (Hartwell & Weinert, 1989), evidences for its existence remain missing.

In contrast to initiation or elongation, the mechanisms that drive DNA replication termination have remained unexplored until recent years (Dewar & Walter, 2017). We now know that the dissociation of the CMG helicase (<u>C</u>DC45, <u>M</u>CM2-7 and <u>G</u>INS) is a key event in termination, which only occurs after the DNA from two converging forks is fully replicated and ligated (Dewar, Budzowska et al., 2015). At the molecular level, replisome disassembly requires MCM7 ubiquitination (Maric, Maculins et al., 2014, Moreno, Bailey et al., 2014), which drives its extraction from chromatin by the VCP segregase and its adaptors UFD1L and NPLOC4 (Maric, Mukherjee et al., 2017, Sonneville, Moreno et al., 2017). In addition, the unloading of the MCM complex in late S-phase is facilitated by MCM-BP (Nishiyama, Frappier et al., 2011). While the E3 ubiquitin ligase complex that ubiquitinates MCM7 upon DNA replication termination has been identified in *C. elegans* and *Xenopus* (CUL2^LRR1^), its mammalian counterpart remains unknown. In this context, we recently reported that USP7 is a replisome-associated deubiquitinase (DUB) that is essential for DNA replication (Lecona, Rodriguez-Acebes et al., 2016). Proteomic analyses revealed that USP7 inhibition increases the ubiquitination of SUMO and SUMOylated factors, which are displaced from replication factories. Interestingly, MCM7 was one of the proteins to show increased ubiquitination upon USP7 inhibition, with the modification occurring at the same region that had been linked to DNA replication termination in yeast (Maric et al., 2017). Moreover, prior work showed that MCM-BP tethers USP7 to MCM proteins, and that lack of USP7 increases the levels of chromatin-bound MCM (Jagannathan, Nguyen et al., 2014). Based on these data, we previously hypothesized that USP7 inhibition could be an approach to mimic DNA replication termination in mammalian cells (Lecona & Fernandez-Capetillo, 2016). Here, we confirmed this hypothesis and used USP7 inhibition as a way to address the potential coordination between the completion of DNA replication and the activation of the mitotic program.

## RESULTS

### USP7 inhibition triggers DNA replication termination

In order to perform a detailed analysis of the effect of USP7 inhibition on cell-cycle progression, human retinal pigmentum epithelial (RPE) cells were synchronized with a double thymidine block and released in the absence or presence of the USP7 inhibitor P22077 (USP7i, hereafter; added 4 hrs after the release). While by 14 hrs control cells had completed an entire cell cycle, the treatment with USP7i led to an immediate arrest in S-phase that could not be overcome, which could be consistent with DNA replication termination (Figure 1A). In support of this, and as previously observed in proteomic analyses (Lecona et al., 2016), USP7 inhibition led to an accumulation of ubiquitinated MCM7 on chromatin, which is a hallmark of replication termination (Maric et al., 2014, Moreno et al., 2014) (Figure 1B). Moreover, MCM7 ubiquitination was also detected in control RPE cells at time-points coincident with mitotic entry and thus with the completion of DNA replication (8-12 hrs). Upon its ubiquitination at the end of DNA replication, MCM7 is extracted from chromatin by the action of the VCP segregase (Maric et al., 2017, Sonneville et al., 2017). Likewise, USP7 inhibition led to an accumulation of the VCP segregase on chromatin concomitant with MCM7 ubiquitination (Figure 1B). Immunofluorescence analysis in U2OS cells confirmed the recruitment of VCP to chromatin upon USP7i treatment (Figure 1C,D).

**Figure 1.**
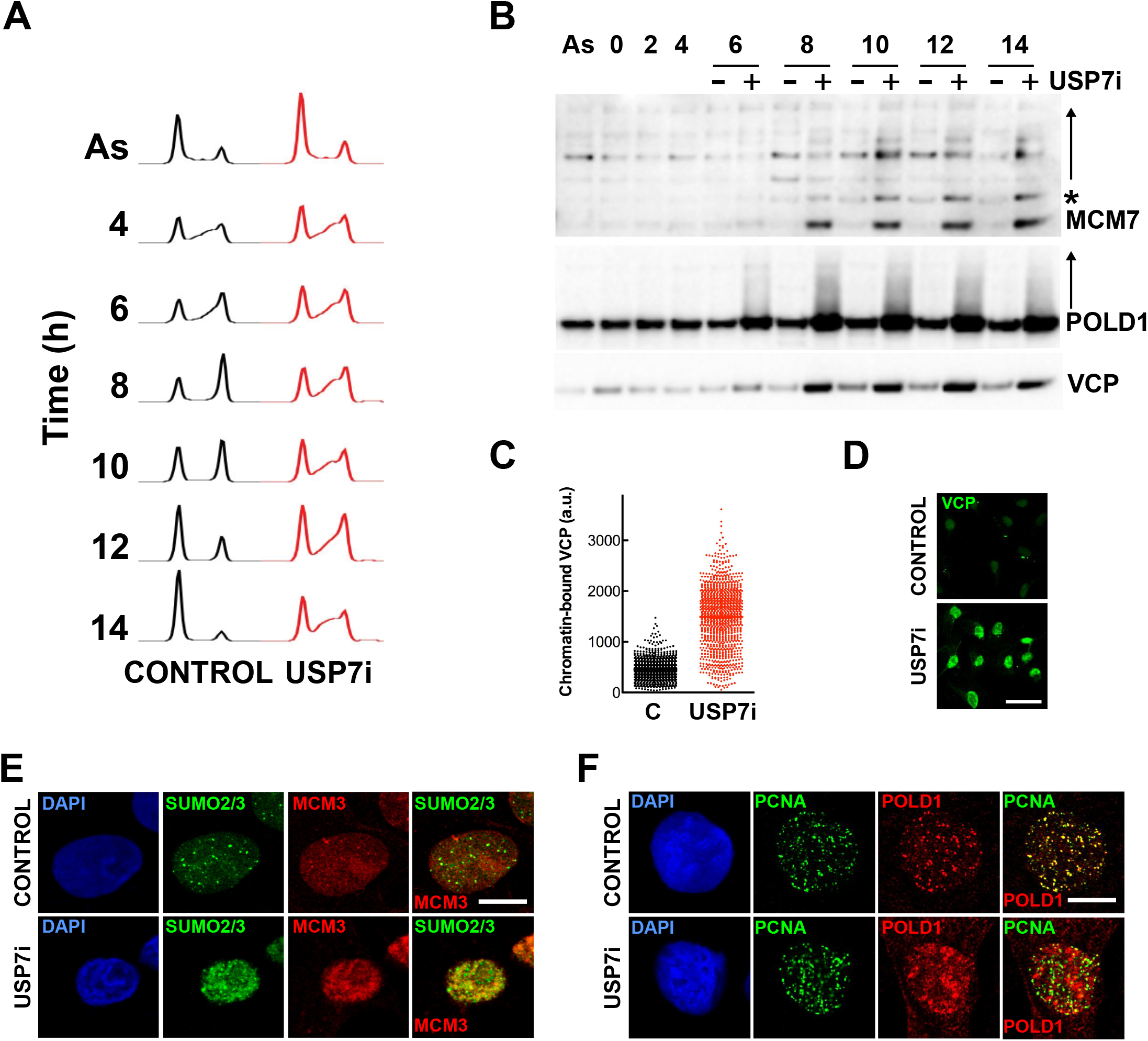
USP7 inhibition mimics DNA replication termination. (A) Cell cycle profile of RPE cells arrested in G1/S and released for the indicated times. The panel on the left shows control cells. On the right cells were treated with 25 μM USP7i 4 hrs after release. Asynchronous cells are shown as control in the top (As). Propidium iodide (PI) was used to measure DNA content. (B) Chromatin-bound levels of MCM7, POLD1 and VCP analysed by Western Blotting (WB) in RPE cells synchronized and treated as in (A). The asterisk indicates the mono-ubiquitinated version of MCM7 and the arrows show the poli-ubiquitination of MCM7 and POLD1. (C-D) Chromatin-bound levels of VCP (green) measured by High-Throughput Microscopy (HTM) in U2OS cells that were either untreated (CONTROL) or after treatment with 50 μM USP7i for 2 hrs. Scale bar, 50 μm. The quantification for this experiment is shown in (C) and a representative image for each condition is shown in (D). (E) Immunofluorescence analysis of chromatin-bound SUMO2/3 (green) and MCM3 (red) levels in U2OS cells that were either untreated (CONTROL) or after treatment with 50 μM USP7i for 4 h. DNA was stained with DAPI (blue). The overlay for SUMO2/3 and MCM3 is shown on the right. Scale bar, 10 μm. (F) Immunofluorescence analysis of chromatin-bound PCNA (green) and POLD1 (red) levels in U2OS cells that were either untreated (CONTROL) or after treatment with 50 μM USP7i for 4 h. DNA was stained with DAPI (blue). The overlay for PCNA and POLD1 is shown on the right. Scale bar, 10 μm.

Similar to the accumulation of VCP, our previous work showed that USP7 inhibition leads to an increase of SUMO2/3 levels on chromatin (Lecona et al., 2016). Interestingly, while SUMO2/3 is normally enriched at replication foci, the increase of chromatin-bound SUMO2/3 triggered by USP7 inhibition occurs away from replisomes, which we suggested was due to a general displacement of SUMOylated replisome factors upon their ubiquitination (Lecona & Fernandez-Capetillo, 2016). Supporting this view, we now show that MCM3, a component of the CMG helicase, accumulates on chromatin together with SUMO2/3 upon exposure to USP7i (Figure 1E). Of note, while much of the focus of DNA replication termination studies has been placed on the role of ubiquitination on the disassembly of the MCM complex (Dewar & Walter, 2017), this mechanism could also apply to other replisome components. In this regard, we also detected an increase in DNA Polymerase delta 1 (POLD1) ubiquitination upon USP7 inhibition in late S-phase (Figure 1B). Immunoprecipitation experiments with histidine-tagged ubiquitin confirmed the ubiquitination of chromatin-bound POLD1 in response to USP7i (Figure S1A). The ubiquitination of POLD1 occurred concomitant to its accumulation on chromatin, but away from sites of DNA replication (as identified by PCNA) (Figure 1F). Of note, other factors such as RPA2 or RFC2 are not delocalized from PCNA foci upon USP7 inhibition (Figure S1B), arguing that the role of USP7 is not a general action on all replisome components. Collectively, these results indicate that USP7 inhibition mimics DNA replication termination, allowing us to explore the potential connection between the end of S-phase and mitotic entry.

**Figure S1.**
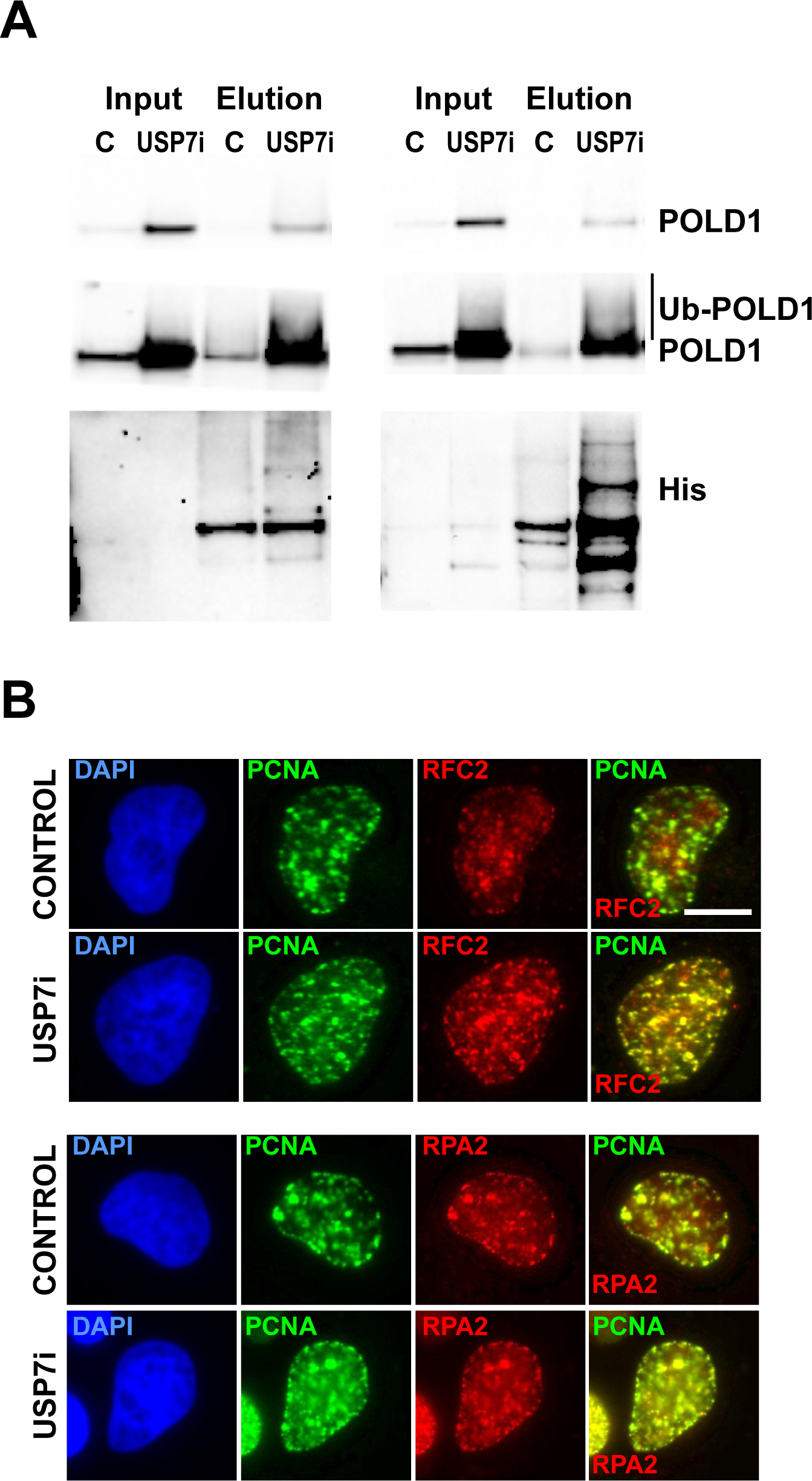
*(Related to Fig. 1)*. (A) Immunoprecipitation with Ni-NTA Agarose of RPE cells transfected with the plasmid pCl-His-Ubiquitin for 48h and treated with 25 μM USP7i or DMSO (Control) for the last 4 h. 2 independent experiments are shown. (B) Immunofluorescence analysis of chromatin-bound PCNA (green) and RFC or RPA2 (red) levels in U2OS cells that were either untreated (CONTROL) or after treatment with 50 μM USP7i for 4 h. DNA was stained with DAPI (blue). The overlay for PCNA and RFC2 or RPA2 is shown on the right. Scale bar, 10 μm.

### USP7 inhibition leads to a premature activation of mitotic kinases

To investigate the effects of USP7 inhibition on the activation of the mitotic program, we first evaluated its impact on the levels of phosphorylated histone H3 Ser 10 (H3S10P), a well-established mitotic mark that is required for chromosome condensation (Wei, Yu et al., 1999), and which was originally discovered in cells undergoing PCC (Ajiro, Nishimoto et al., 1983). Despite the immediate arrest of DNA replication that is observed upon a treatment with USP7 inhibitors, exposure of human HCT116 cells to these agents led to a significant increase in H3S10 phosphorylation (Figure 2A). Surprisingly, flow cytometry analyses revealed that UPS7 inhibition led to a global accumulation of H3S10P throughout the cell cycle (Figure 2B), rather than being restricted to mitotic cells. Similar observations were done using MPM-2, a monoclonal antibody raised against mitosis-specific phosphorylation events (Davis, Tsao et al., 1983), which also marks sites of chromosome condensation (Rieder & Cole, 1998) (Figure 2A,B). Importantly, these effects were observed in all cell lines tested including RPE and U2OS (Figure S2A,B). Moreover, they were recapitulated with an independent USP7 inhibitor (Ritorto, Ewan et al., 2014) (Figure S2C), and by RNA interference-mediated depletion of USP7 (Figure 2C-E), confirming the selectivity of the effect.

**Figure 2.**
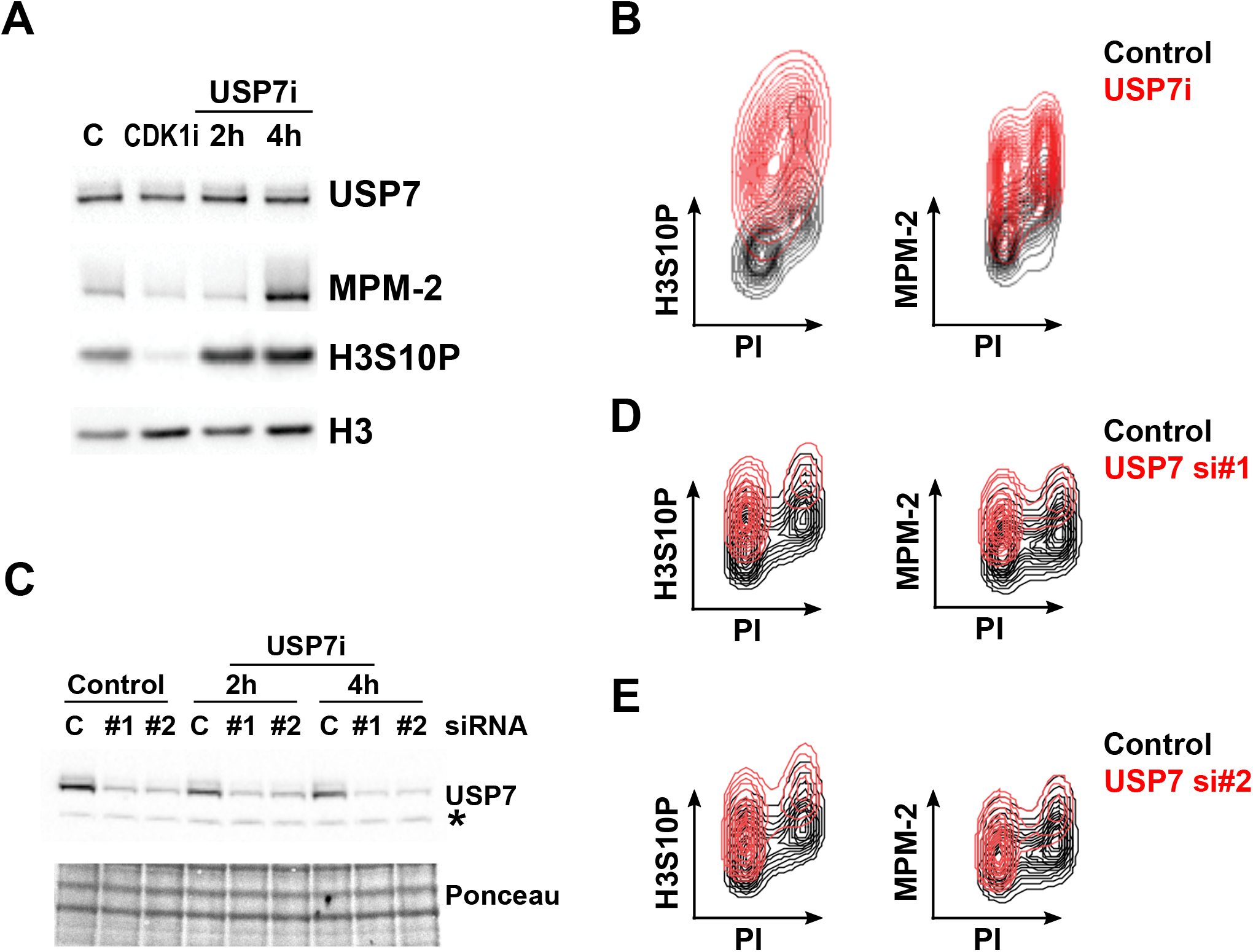
USP7 restricts the activity of mitotic kinases. (A) WB showing the levels of USP7, MPM-2, histone H3S10P and histone H3 in whole nuclear extracts of HCT116 cells treated with 50 μM USP7i for 2-4 h, 10 μM of the CDK1 inhibitor Ro3306 for 8 hrs (CDK1i, hereafter) or DMSO as a control (C). (B) Flow cytometry data illustrating the levels of histone H3S10P (left) and MPM-2 (right) in HCT116 cells that were either untreated (CONTROL, black) or after treatment with 50 μM USP7i for 8 hrs (red). DNA content was measured with Propidium Iodide (PI). (C) WB showing the levels of USP7 in whole cell extracts from RPE cells transfected with a non-specific siRNA or 2-independent siRNAs against USP7. Cells were treated with 25 μM USP7i for 2-4 hrs or DMSO as a control (CONTROL). Ponceau staining is shown as loading control and an asterisk indicates a non-specific band. (D-E) Flow cytometry data illustrating the levels of histone H3S10P and MPM-2 in RPE cells transfected with two different specific siRNA against USP7 (red) or with a non-specific siRNA (Control, black), corresponding to the Western Blot in (C). DNA content was measured with Propidium Iodide (PI). The experiments were repeated 3-5 times and one representative result is shown. See also Figure S1.

**Figure S2.**
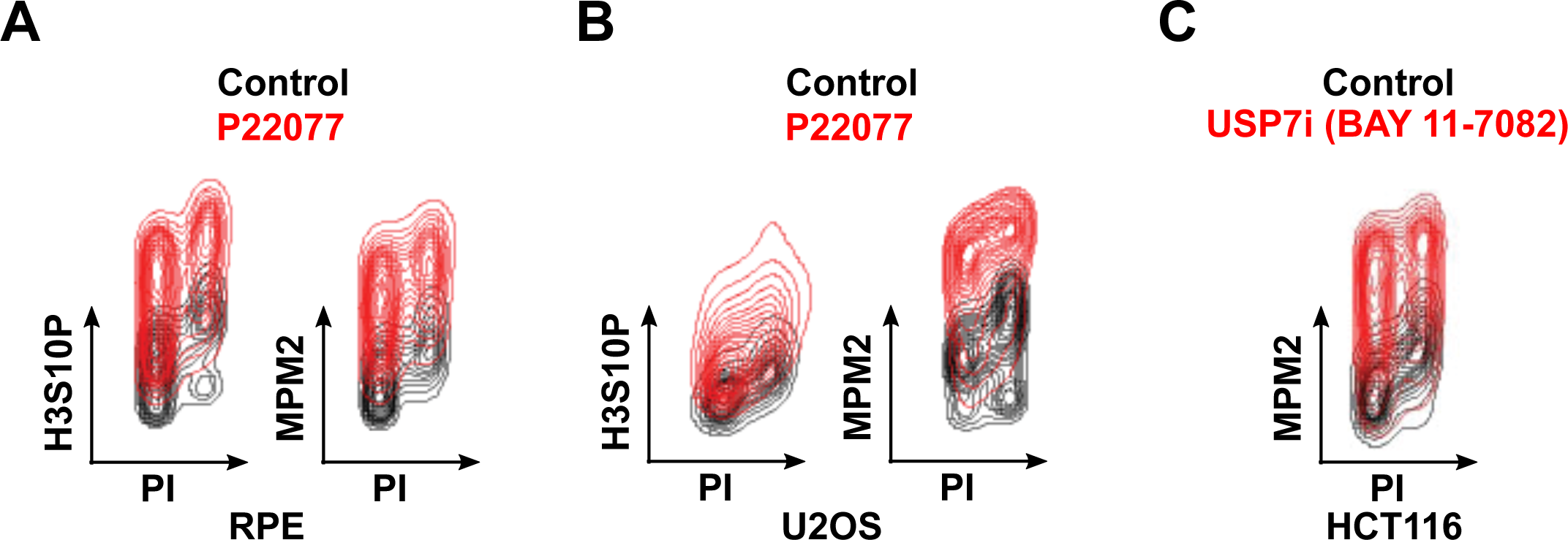
*(Related to Fig. 2)*. (A-B) Flow cytometry data illustrating the levels of histone H3S10P (left) and MPM-2 (right) in RPE (A) and U2OS (B) cells that were either untreated (CONTROL, black) or after a treatment with 25 or 50 μM USP7i for 4 hrs (red), respectively. DNA content was measured with PI. The experiments were repeated 2-3 times and one representative result is shown. (C) Flow cytometry data illustrating the levels of MPM-2 in RPE cells that were either untreated (CONTROL, black) or after treatment with 25 μM of an independent USP7 inhibitor (BAY 11-7082) for 2 hrs (red).

Mitotic entry is primarily driven by the activation of CDK1 upon the nuclear translocation of cyclin B1 (CCNB1), and the removal of inhibitory phosphorylations at Thr-14 and Tyr-15 by the CDC25 family of phosphatases (reviewed in (Rhind & Russell, 2012)). Consistent with the global activation of mitotic kinases observed by flow cytometry, immunofluorescence analyses revealed a generalized accumulation of nuclear CCNB1 in RPE cells treated with USP7i, which was phosphorylated at Ser-126, a CDK1-dependent site (Li, Meyer et al., 1997) (Figure 3A,B). In addition, WB confirmed an increase of other CDK1-dependent phosphorylation events such as LAMIN A/C or MPM-2, together with a decrease in CDK1-Tyr15 phosphorylation, all of which are indicative of CDK1 activation (Figure 3C). Supporting this view, CDK1 inhibition rescued the generalized increase in H3S10 phosphorylation and MPM-2 reactivity that is observed upon USP7 inhibition (Figure 3D,E). Moreover, we tested the effect of USP7 inhibitors in mouse embryonic fibroblasts (MEF) lacking CDK2, CDK4 and CDK6 (CDK2/4/6 KO), in which CDK1 drives all cell cycle transitions by interacting with the corresponding cyclins (Santamaria, Barriere et al., 2007). MPM-2 and H3S10 phosphorylation presented a similar increase in Wild Type (WT) and CDK2/4/6 KO MEFs treated with USP7 throughout the cell cycle, further indicating that this activity is mediated by CDK1 (Figure 3F,G).

**Figure 3.**
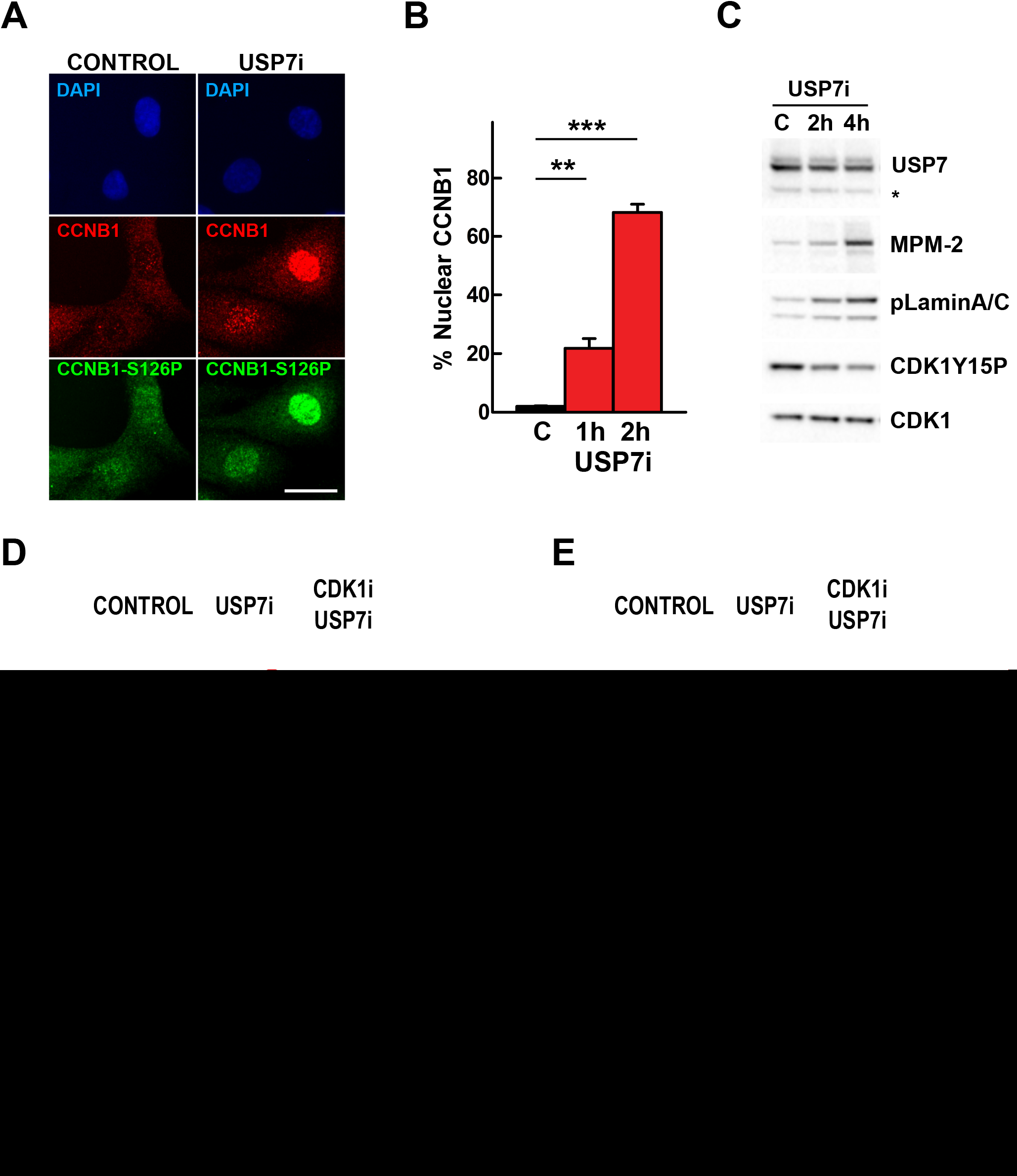
USP7 inhibition activates CDK1. (A-B) Immunofluorescence of CCNB1 (red) and CCNB1S126P (green) in RPE cells that were either untreated (CONTROL) or after treatment with 25 μM USP7i for 2 h. DNA was stained with DAPI (blue). Scale bar, 30 μm. (B) HTM-mediated quantification of the percentage of nuclear CCNB1 from RPE treated as in (A). The graph provides the mean and s.d. from three independent experiments (**p<0.01, ***p<0.005, t-test). (C) WB showing the levels of USP7, MPM-2, pLaminA/C, CDK1Y15P, and CDK1 in whole cell extracts of RPE cells treated with 25 μM USP7i for 2-4 h. (D-E) Flow cytometry data illustrating the levels of MPM-2 (D) and H3S10P (E) in RPE cells that were either untreated (CONTROL, black), treated with 25 μM USP7i for 4 hrs (red) or with 10 μM CDK1i for 8 hrs followed by incubation with 25 μM USP7i and 10 μM CDK1i for 4 additional hours (blue). DNA content was measured with PI. One representative experiment is shown, out of five. (F-G) Flow cytometry data illustrating the levels of MPM-2 (F) and H3S10P (G) in WT and CDK2/4/6 KO MEFs that were either untreated (CONTROL, black) or treated with 50 μM USP7i for 4 hrs (red). The experiment was performed twice using two independent WT and CDK2/4/6 KO MEF pairs and one representative result is shown.

### CDK1 and CDC25A contribute to the toxicity of USP7 inhibitors

Finally, we evaluated if the premature activation of CDK1 plays a role on the toxicity of USP7 inhibitors, which are being developed as anticancer agents mostly based on their capacity to increase P53 levels (Li, Chen et al., 2002). As mentioned above, a premature activation of CDK1 in S-phase cells leads to PCC, abnormal chromosome segregation and ultimately cell death (Aarts et al., 2012, Duda et al., 2016). Likewise, metaphase spreads prepared from RPE cells treated with USP7i revealed figures consistent with PCC and a significant increase in aberrant metaphases (Figure 4A,B). Importantly, CDK1 inhibition significantly reduced the percentage of apoptotic cells induced by USP7i (Figure 4C,D). We recently showed that the toxicity of inhibitors of the ATR kinase is also mediated by an increase in CDK1 activity, and which is alleviated by a deficiency in the CDK1-activating CDC25A phosphatase (Ruiz et al., 2016). Similarly, CDC25A-deficient mouse embryonic stem cells were significantly resistant to USP7i-induced cell death (Figure 4E,F). Hence, and regardless of the effect of USP7 inhibitors on P53, our data show that the toxicity of these compounds is due, at least in part, by a premature activation of CDK1.

**Figure 4.**
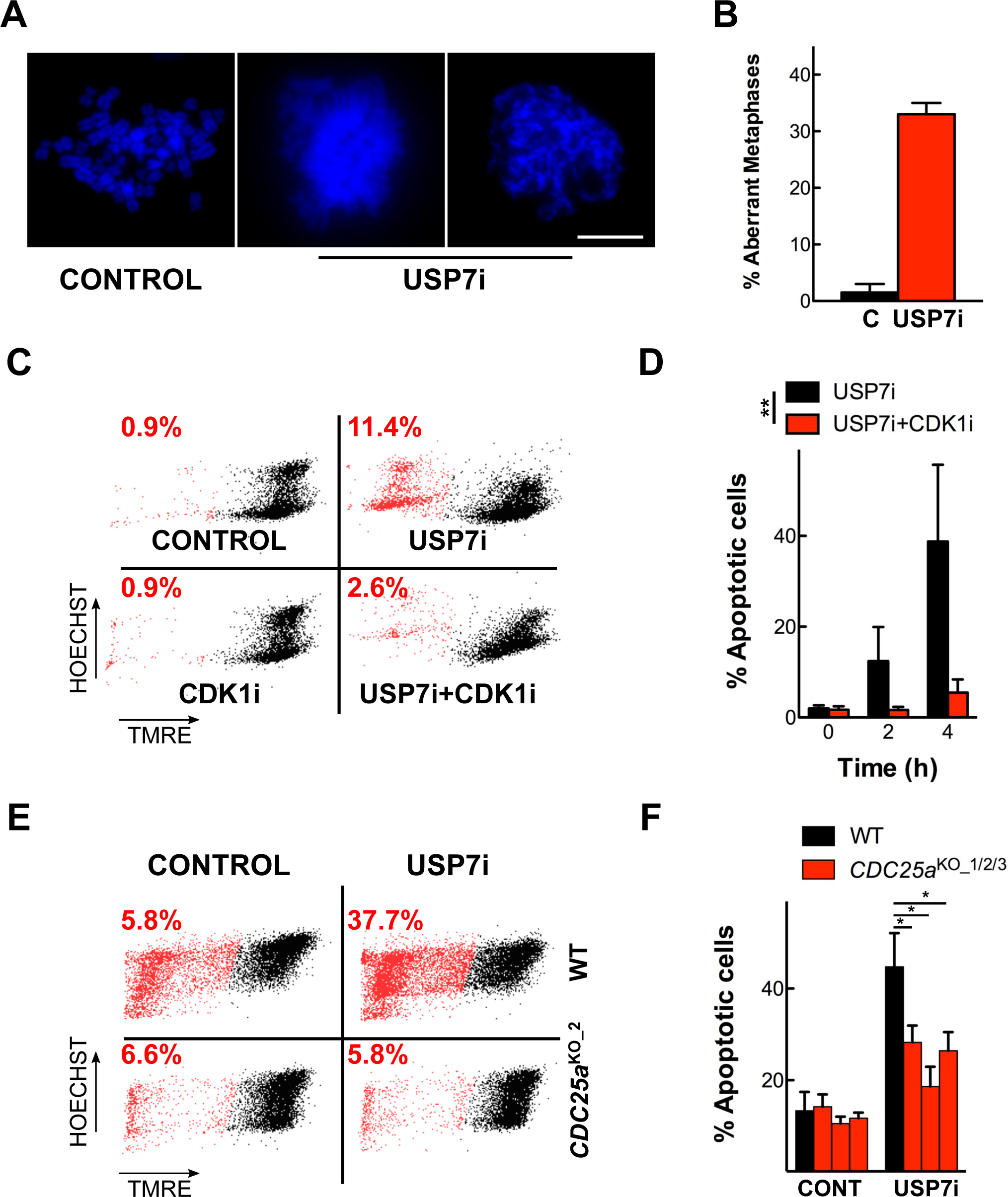
USP7 inhibition induces CDK1-dependent cell death. (A-B) Metaphase spreads of RPE cells arrested in S phase with Thymidine, released for 2 hrs and then treated with DMSO (CONTROL) or 25 μM USP7i for 2 h. Scale bar, 10 μm. The percentage of aberrantly condensed metaphases was quantified from 2 independent experiments (B). Bars indicate s.d. (C-D) Flow cytometry analysis of apoptosis in RPE cells after treatment with DMSO (CONTROL), 25 μM USP7i, 10 μM CDK1i, or a combined treatment of 25 μM USP7i and 10 μM CDK1i for 4 h. DNA content was evaluated with Hoechst, and the mitochondrial membrane potential was measured by TMRE. The percentage of apoptotic cells (TMRE negative, red) is indicated for each condition from a representative experiment. The experiment was repeated 3 times and the quantification is shown in (D) (**p<0.01, ANOVA). (E-F) Flow cytometry analysis of apoptosis in WT and *CDC25a*^KO^ mESC after treatment with DMSO as a control (CONTROL) or with 25 μM USP7i for 16 h. DNA content was evaluated with Hoechst, and the mitochondrial membrane potential was measured by TMRE. The percentage of apoptotic cells (TMRE negative, red) is indicated for each condition from a representative experiment. The experiment was repeated 3 times using 3 independent CDC25A-deficient mESC lines. The quantification is shown in (F). (*p<0.05, t-test).

## DISCUSSION

While the cell cycle has been intensively studied, our understanding of how or when mitosis is initiated remains incomplete. A passive model would be that a gradual build-up of CDK1 activity in G2 that, at some point, reaches a threshold sufficient to trigger mitotic entry. Alternatively, or in addition to this model, it has been long speculated that an S/M checkpoint constantly prevents mitotic entry before DNA replication is completed (Elledge, 1996, Enoch & Nurse, 1991, Hartwell & Weinert, 1989). In this model, G2 could *de facto* be considered as a “very-late S phase”, where cells would have completed the replication most of their genome, with some difficult parts such as repetitive sequences or heterochromatin regions left to be copied. Interestingly, recent works have detected evidences of DNA replication persisting in mitosis, which could be exacerbated by conditions that perturb DNA replication (Eykelenboom, Harte et al., 2013, Minocherhomji, Ying et al., 2015).

The S/M checkpoint model also implies that replisomes will somehow elicit a “No-Mitosis” signal, that will disappear once replication is finished allowing for the progression into mitosis. Supporting this view, previous work revealed that the activation of mitotic kinases is coincident with the disappearance of PCNA foci (Akopyan, Silva Cascales et al., 2014). This concept would be analogous to the <u>S</u>pindle <u>A</u>ssembly <u>C</u>heckpoint (SAC), where every unaligned chromosome actively signals through the SAC to prevent the onset of anaphase (Musacchio, 2015). In the S/M checkpoint, ongoing replisomes would be the signal that prevents CDK1 activation. One possibility could be that the single-stranded DNA generated at moving forks suffices to activate the ATR-dependent checkpoint, which limits CDK1 activity. Accordingly, ATR inhibition forces premature mitotic entry in cells that have unfinished DNA replication (Eykelenboom et al., 2013, Ruiz et al., 2016). In addition to kinases, our work here indicates that ubiquitin-based signaling networks also play a role in coordinating S and M phases, as the inhibition of a replisome-associated DUB (USP7) leads to the activation of CDK1. How an increase in ubiquitin modifications leads to CDK1 activation remains unclear at this point. Intriguingly, CDK1 is among the factors that we previously found to be ubiquitinated in response to USP7 inhibition (Lecona et al., 2016), and recent work has revealed that SUMOylation is inhibitory for CDK1 function (Xiao, Lucas et al., 2016). One possibility could thus be that SUMO and ubiquitin play opposing roles in regulating CDK1 activity, although whether this occurs and how remains to be clarified.

Finally, our work also has important implications for the development of USP7 inhibitors as anticancer agents, an area in which efforts have recently intensified (reviewed in (Zhang & Sidhu, 2018)). The currently accepted mechanism of action is the deubiquitination and stabilization of P53 due to the degradation of MDM2, the main E3 ligase targeting P53 for degradation by the proteasome (Li et al., 2002). However, our previous work showing USP7 inhibition blocks DNA replication independently of p53 (Lecona et al., 2016), together with the current data indicating that the toxicity of these compounds is alleviated by CDK1 inhibition or CDC25A deficiency, indicate that premature mitotic entry is likely to play an important role on the toxic effects of USP7 inhibitors. This information should be incorporated into the rationale for using these compounds, first because it indicates that p53-deficient cancer cells will also respond to the treatment, but also since it suggests that USP7 inhibitors are likely to synergize with other agents that promote mitotic entry. Besides its usefulness for cancer therapy, the work presented here provides direct evidence for the coupling of DNA replication termination with mitotic entry and identifies USP7 as a coordinator of this transition.

## AUTHOR CONTRIBUTIONS

A.G. and E.L. designed and participated in most of the experiments of this study. P.V. and P.U. helped with experiments from Figure 1. V.L. helped with immunofluorescence and metaphase analyses. O.F. and E.L. coordinated the study and wrote the MS.

## ACKNOWLEDGEMENTS

Research was funded by Fundación Botín, Banco Santander, through its Santander Universities Global Division and by grants from the Spanish Ministry of Economy and Competitiveness (MINECO) (SAF2014- 57791-REDC), the Howard Hughes Medical Institute and the European Research Council (ERC-617840) to OF; a grant from MINECO (BFU2014-55168-JIN) that is co-funded by European Regional Development Funds (FEDER) to EL; and a PhD fellowship from MINECO to AG (BES-2015-075758). The authors declare no competing financial interests.

## MATERIALS AND METHODS

### Cell lines

HCT116, U2OS and RPE cells (ATCC) were grown in DMEM with 10% FBS, penicillin (100 IU/ml), streptomycin (100 mg/ml) and glutamine (300 mg/ml). MEFs were grown in 15% FBS, penicillin (100 IU/ml), streptomycin (100 mg/ml) and glutamine (300 mg/ml). mESCs were grown on a feeder layer of inactivated MEF with DMEM (high glucose) supplemented with 15% knockout serum replacement (Invitrogen), LIF (1000 U/ml), 0.1 mM non-essential aminoacids, 1% glutamax and 55 μM β-mercaptoethanol.

### Treatments

USP7 inhibitors P22077 (Merck Millipore), BAY 11-7082 (Santa Cruz) and the CDK1 inhibitor Ro3306 (Sigma) were dissolved in DMSO. Cells were incubated for the indicated times and doses with each inhibitor or an equivalent amount of DMSO.

### Cell Synchronization

RPE cells were synchronized using a double thymidine block. Cells were incubated in the presence of 1 mM Thymidine for 16hrs at 37°C. Then, cells were washed once in PBS and released in DMEM for 8hrs at 37°C. The culture medium was replaced with DMEM containing 1 mM Thymidine and cells were incubated for 16hrs at 37°C. Again, cells were washed once and released in DMEM.

### Immunofluorescence and High Throughput Microscopy

For immunofluorescence of MCM3, SUMO2/3, VCP, PCNA, POLD1, RFC2 and RPA2 soluble proteins were pre-extracted by mild detergent permeabilization with CSKI buffer (10 mM Pipes, pH 6.8, 100 mM NaCl, 300 mM sucrose, 3 mM MgCl2, 1 mM EGTA, and 0.5% Triton X-100) before fixation. For HTM, cells were grown on μCLEAR bottom 96-well plates (Greiner Bio-One) and immunofluorescence was performed using standard procedures. Analysis of DNA Replication by EdU incorporation was done using Click-It (Invitrogen) following manufacturers’ instructions. In all cases, images were automatically acquired from each well using an Opera High-Content Screening System (Perkin Elmer). A 20x or 40x magnification lens was used and images were taken at non-saturating conditions. Images were segmented using DAPI signals to generate masks matching cell nuclei from which the mean signals for the rest of the stainings were calculated.

### Protein extracts and Cell Fractionation

Whole cell extracts were prepared by lysing cells in 50 mM Tris, pH 7.5, 8 M Urea, and 1% Chaps. Cytosolic and nuclear extracts were prepared as described (Lecona, Barrasa et al., 2008). Cells were scraped in cold PBS and washed twice with PBS. Cells pellets were resuspended in ice-cold hypotonic lysis buffer (10 mM HEPES, pH 7.9, 10mM KCl, 0.1 mM EDTA containing protease and phosphatase inhibitors), incubated on ice for 10 min, and then Nonidet P-P40 was added to a final concentration of 0.1%. After 3 min at room temperature, cells were vortexed and the cytosolic fraction was obtained by centrifugation at 2500 *g* for 5 min. Nuclei were washed once in hypotonic lysis buffer and then resuspended in hypertonic extraction buffer (20 mM HEPES, pH 7.9, 0.4 M NaCl, 1 mM EDTA containing protease and phosphatase inhibitors). After 1 hr shaking at 4°C, the extract was centrifuged for 5 min at 16000 *g*. The nuclear extract is collected and the chromatin fraction was then extracted in 50 mM Tris, pH 7.5, 8 M Urea, and 1% Chaps, shaking at 4°C for 30 min. Chromatin fraction was obtained after centrifugation for 5 min at 16000 *g*. Protein concentration was determined using the BioRad Protein Assay.

### Immunoprecipitation of His-ubiquitin

Transfection of RPE cells with the plasmid pCl-His-hUb (Addgene, #31815) was carried out using Lipofectamine 2000 (Invitrogen) according to the manufacturer’s instructions. After cell fractionation as mentioned before, the chromatin fraction was diluted in 8 M urea/100 mM phosphate/ 500 mM NaCl/30 mM imidazole/0.5% NP40, and then incubated in Ni-NTA Agarose (Qiagen), for 1.5 h, rotating at RT. The beads were washed with 8 M urea/100 mM phosphate/ 500 mM NaCl/30 mM imidazole/0.5% NP40, and eluted with 8 M urea/100 mM phosphate/400 mM imidazole.

### Flow Cytometry

For the analysis of H3S10P and MPM-2, cells were trypsinized, washed with cold PBS once and fixed in suspension by the addition of cold 70% ethanol. Cells were incubated for 30 min on ice and maintained at −20°C or processed immediately after fixation. Cells were centrifuged at 4000 rpm for 5 min and then incubated with the specific antibodies in PBS/0.05% Tween20/ 3% BSA for 2 hrs at RT. Cells were washed in PBS/0.05% Tween20 and incubated with secondary antibodies in PBS/0.05% Tween20/ 3% BSA for 1 hr at RT. Finally, cells were washed with PBS/0.05% Tween20 and resuspended in PBS containing 100 μg/ml RNase. DNA content was visualized incubating the cells with 100 μg/ml PI. For the analysis of apoptosis, cells were trypsinized, centrifuged and resuspended at 10^6^ cells/ml in cell culture medium containing 10 μg/ml Hoechst and 40nM TMRE. Then, cells were incubated for 30 min at 37°C. All samples were analysed in a BD LSRFortessa cell analyser. The results were analysed using the FlowJo software (FlowJo, LLC).

### RNA interference

Transfection of RPE cells with control (Qiagen; Cat#1027281) or USP7 targeting siRNAs (USP7 siRNA #1 (Cat#SI00052283) and #2 (Cat#SI00052290)) was carried out using Lipofectamine RNAimax (Invitrogen) according to the manufacturer’s instructions.

### Metaphases

Cultured cells were arrested at mitosis with 100 ng/ml Colcemid (GIBCO/BRL) and after 2 hours were collected and resuspended in a hypotonic solution (0.075 mM of KCl). Cells were incubated 20 minutes at 37°C and fixed with Carnoy´s buffer (methanol-glacial acetic acid (3:1)). To obtain metaphase spreads cells were dropped on slides and stained with Giemsa.

### Data analysis

Data were represented with the use of the Prism software (GraphPad Software). The statistical analysis was also carried out with Prism, using the specified statistical comparison where indicated in the Figure Legends.

### Antibodies

The following antibodies were used in this study: H3S10P (Merck Millipore, 06-570), H3 (Abcam, ab10799), MPM-2 (Merck Millipore, 05-368), USP7 (Bethyl, A300- 033A), POLD1 (Santa Cruz, sc-10784), MCM7 (LSBio, LS-C331288), MCM3 (kind gift of Dr. Juan Mendez), SUMO2/3 (MBL, M114-3), VCP (GeneTex, GTX101089), PCNA (Santa Cruz, sc-56; Merck Millipore, 07-2162), RFC2 (GeneTex, GTX105087), POLD1 (Santa Cruz, sc-373731, sc-373731), RPA2 (Cell Signaling, 2208), CCNB1 (Santa Cruz, sc-245), CCNB1-S126P (Abcam, ab55184), Lamin AC-S22P (Cell Signaling, 2026), CDK1-Y15P (Santa Cruz, sc-7989), CDK1 (Merck Millipore, 06-923), goat anti-rabbit IgG (H+L)-HRP (ThermoFisher, 31460), goat anti-mouse IgG (H+L)-HRP (ThermoFisher, 31430), Alexa Fluor 488 anti-mouse (ThermoFisher, A11001), Alexa Fluor 488 anti-rabbit (ThermoFisher, A21441), Alexa Fluor 594 anti-mouse (ThermoFisher, A11005) and Alexa Fluor 647 antirabbit (ThermoFisher, A21443).

